# Somatostatin neurons of the bed nucleus of stria terminalis enhance associative fear memory consolidation in mice

**DOI:** 10.1101/2020.05.22.103481

**Authors:** Biborka Bruzsik, Laszlo Biro, Dora Zelena, Eszter Sipos, Huba Szebik, Klara Rebeka Sarosdi, Orsolya Horvath, Imre Farkas, Veronika Csillag, Cintia Klaudia Finszter, Eva Mikics, Mate Toth

## Abstract

Excessive fear learning and extinction-resistant fear memories are core symptoms of anxiety and trauma-related disorders. Despite significant evidence from clinical studies reporting hyperactivity of the bed nucleus of stria terminalis (BNST) under these conditions, the role of BNST in fear learning and expression is still not clarified. Here, we tested how BNST modulates fear learning in mice using a chemogenetic approach. Activation of GABAergic neurons of BNST during fear acquisition, more specifically the consolidation phase, resulted in enhanced cued fear recall. Importantly, BNST activation had no acute impact on fear expression during conditioning or recalls, but it enhanced cued fear recall subsequently, potentially via altered activity of downstream regions as indicated by c-Fos. Enhanced fear memory consolidation could be replicated by selectively activating somatostatin neurons (but not corticotropin releasing factor neurons), suggesting significant modulation of fear memory strength by specific circuits of BNST.

## Introduction

Forming memories of negative events are essential for the survival of the individual, however, excessive fear learning and extinction-resistant memory formation results in maladaptive, inflexible phenotype or symptoms present in anxiety and trauma-related disorders, i.e. posttraumatic stress disorder (PTSD) (Duits *et al*, 2015; Singewald *et al*, 2015). The extended amygdala, including the bed nucleus of stria terminalis (BNST), shows activation during acute threats and sustained anxiety-like states in healthy populations, which becomes elevated in patients with anxiety disorders (Avery *et al*, 2016; Brinkmann *et al*, 2017; Klumpers *et al*, 2017). Early studies investigating the rodent BNST observed functional division between the amygdala and BNST, which was supported by human fMRI data. According to this model, amygdala mediates imminent phasic ‘fear-like’ states, whereas BNST mediates more diffuse unconditioned ‘anxiety-like’ states (Daniel and Rainnie, 2016; Davis *et al*, 2010; Goode *et al*, 2019). Recently, accumulating evidence showed that both amygdala and BNST are recruited under both phasic and sustained (or conditioned and unconditioned) fear-like states in humans and primates (Gungor and Paré, 2016; Shackman and Fox, 2016). *In vivo* electrophysiological recordings in rodents also demonstrated that BNST neurons are recruited during fear acquisition and conditioned stimulus dependent fear recall, when stimuli are not diffuse as a threatening context (Bjorni *et al*, 2020; Daldrup *et al*, 2016; Haufler *et al*, 2013; Jennings *et al*, 2013), indicating that the adjustment of the above model is timely. Recent models reconcile the controversies by emphasizing the dimensionality of fearful stimuli, i.e. the recruitment of BNST is gradiently increased by unpredictability (Goode and Maren, 2017; Goode *et al*, 2019; Gungor *et al*, 2016), and hence, it has a potential role in fear generalization (De Bundel *et al*, 2016; Duvarci *et al*, 2009).

Our study investigated how BNST hyperactivation (as recognized in anxiety disorders) modulates fear learning characteristics in a Pavlovian fear conditioning paradigm that is suitable for translational studies by targeting conserved mechanisms (Deslauriers *et al*, 2018; Flandreau and Toth, 2018). We investigated different phases of fear learning, i.e. fear acquisition, consolidation, and recall evoked by contextual and conditioned stimuli (CS), and their generalization to safety context and cues. Additionally, we aimed to dissect how genetically defined cell populations contribute to these processes, considering significant heterogeneity and complexity of BNST circuits (Daniel *et al*, 2016; Gungor *et al*, 2016; Jennings *et al*, 2013; Kim *et al*, 2013). To address these questions, we applied c-Fos immunohistochemistry that revealed significant activation of BNST during fear acquisition, but not during CS-dependent fear recall. In accordance, tonic chemogenetic activation of the major cell type of BNST (i.e. vesicular GABA transporter positive neurons, BNST^vGAT^) during fear conditioning resulted in enhanced CS-dependent fear recall (freezing), which was replicated when BNST^vGAT^ neurons were activated during fear memory consolidation. In line with our c-Fos data, chemogenetic activation during fear expression (recall) did not alter freezing levels. Finally, we applied chemogenetic activation and inhibition of somatostatin (BNST^SOM^) and corticotropin releasing factor expressing (BNST^CRF^) neurons during fear memory consolidation based on their opposing role in the central amygdala, i.e. driving active and passive fear responses, respectively (Fadok *et al*, 2017; Sanford *et al*, 2017). BNST^SOM^, but not BNST^CRF^, activation resulted in markedly enhanced CS-induced fear recall. These data suggest both functional similarities and differences between amygdala and BNST in the regulation of the fear response: BNST does not drive acute fear response, but it modulates memory strength potentially via plasticity changes in the fear circuitry.

## Results and Discussion

### BNST is activated during cued fear acquisition, but not during recall

To visualize the recruitment of BNST during fear learning and expression, we mapped neuronal activity during fear acquisition (i.e. conditioning) and CS-induced fear recall in adult male C57Bl/6J mice by means of c-Fos immunohistochemistry. Mice were sacrificed for c-Fos staining 90 min after either fear conditioning or CS-dependent fear recall 2 days later, where control mice were exposed to same stimuli (contextual and auditory) without footshocks. Fear conditioning induced significant c-Fos expression in all BNST subregions, particularly in the anteromedial and posterior regions (Figure 1A and D; all regions: F1,18>5.55, p<0.029), confirming previous electrophysiological data showing excitation of BNST neurons during fear learning (Bjorni *et al*, 2020; Haufler *et al*, 2013). In contrast, fear recall was not associated with BNST activation (Figure 1B and E; all tests: F_1,19_>1.008, p>0.327). These data suggest that BNST is recruited during cued fear conditioning, but not when fear is recalled. Accordingly, we aimed to test how this activity contributes to fear learning.

**Figure 1.**
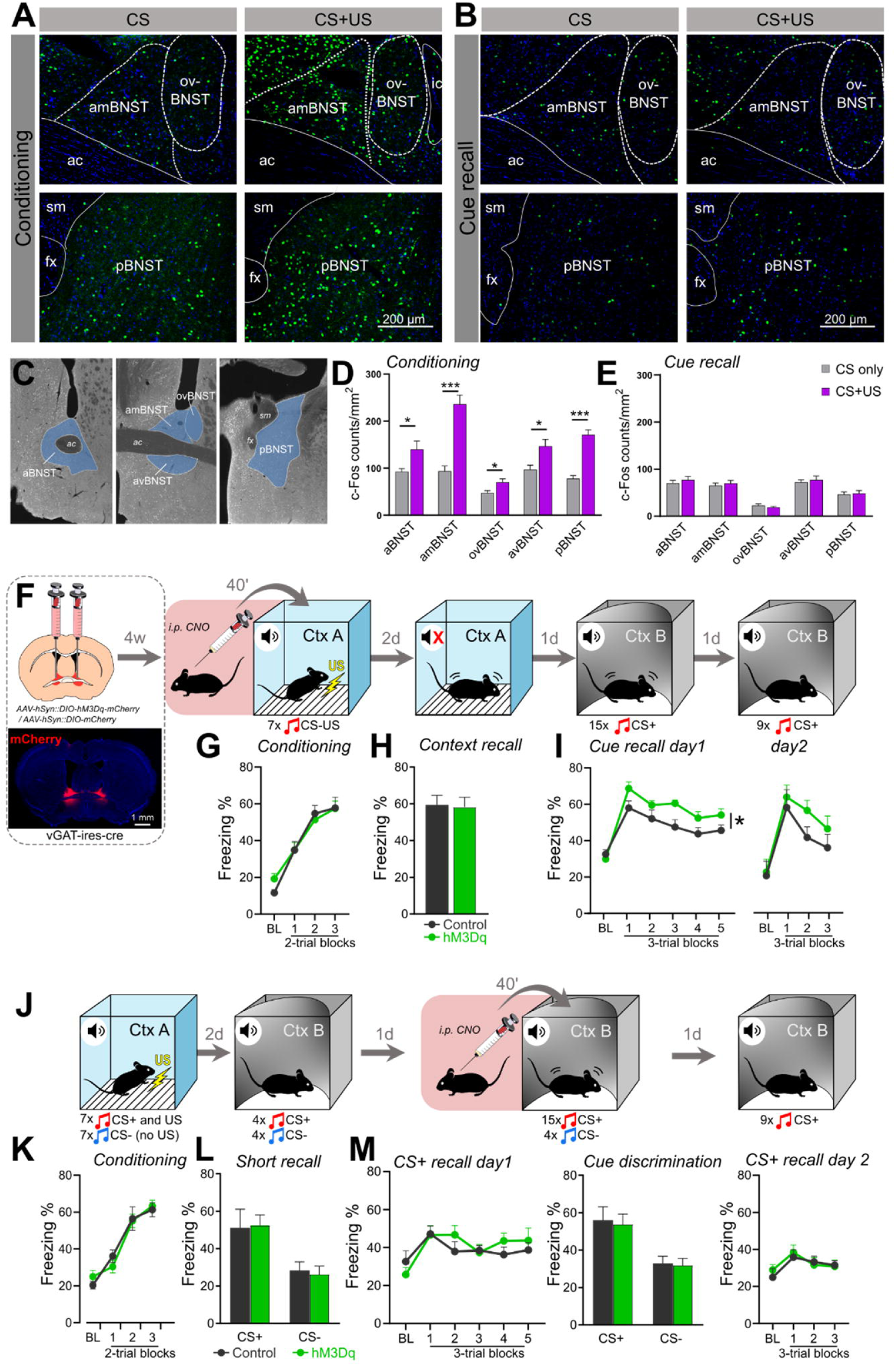
BNST facilitates fear learning, but not fear recall. Representative fluorescent images showing c-Fos immunostaining of BNST subregions during auditory fear conditioning **(A)** and auditory fear recall **(B). (C)** Illustration of investigated BNST subregions for c-Fos quantification. Control mice (gray bars) were exposed to the auditory conditioned stimulus (CS) only without footshocks, whereas conditioned animals received 7 pairings of shock-cue (CS+US) pairings (purple bars). **(D)** Fear conditioning significantly increased c-Fos expression of all BNST subregions (n=10 / groups), in contrast to cued fear recall where no changes in c-Fos expression could be detected **(E).** (n= 10-11 / groups) **(F)** Experimental design for chemogenetic modulation of BNST during fear conditioning. Left dashed-line box illustrates the injection of AAVs encoding hM3Dq-mCherry or mCherry into vGAT-ires-cre mice 4 weeks before behavioral testing, with a representative coronal section of BNST depicting hM3Dq-mCherry expression. Chemogenetic activation did no affect freezing response during fear conditioning **(G)** or contextual fear recall **(H)**, however, enhanced cued fear recall during the first recall session **(I).** (Control: n=8, hM3Dq: n=9) **(J)** Experimental design for chemogenetic activation of BNST during cued fear recall with conditioned (CS+) and safety cue (CS-) presentations. **(K)** Freezing response of hM3Dq and control groups were similar during all testing phase, i.e. conditioning **(L)**, short CS+ and CS-recall **(L)**, CS+ and CS-induced fear recall and their discrimination index **(M).** (Control: n=8, hM3Dq: n=13) *On freezing time curves, each major tick depicts a 150 s block, starting with a pre-tone baseline period (BL). All data are represented as means ± s.e.m. Asterisks represent main effect of ANOVA: *p<0.05; **p<0.01, ***p<0.001. Abbreviations: aBNST: anterior BNST, ac: anterior commissure, amBNST: anteromedial BNST, avBNST: anteroventral BNST, CS/CS+: conditioned stimulus, CS-: stimulus not paired with shock=‘safety signal’, Ctx A: context A (conditioned context), Ctx B: context B (safe context), fx: jornix, ic: internal capsule, ovBNST: oval nucleus of the BNST, pBNST: posterior BNST, sm: stria medullaris; US: unconditioned stimulus.*

### Chemogenetic activation of BNST facilitates fear learning, but not fear expression

We expressed stimulatory hM3Dq ‘Designer Receptors Exclusively Activated by Designer Drugs’ (DREADD) in the BNST of adult male vGAT-ires-cre mice to target the major neuronal population of BNST, i.e. GABAergic neurons (Figure 1 – figure supplement 1). Control animals were injected with viral vector carrying only mCherry fluorophore protein without active hM3Dq receptor. Consistently with previous studies (Mazzone *et al*, 2018), whole-cell patch-clamp recordings confirmed that clozapine-N-oxide (CNO) activated hM3Dq receptors indicated by depolarized resting membrane potential (4.18 ± 0.60 mV vs. – 0.54 ± 1.11 mV in controls; F_1,7_=17.319, p=0.004; t=9.286, p<0.001 compared to baseline) and increased firing rate of hM3Dq-expressing BNST^vGAT^ cells (30 pA pulse: from 0.9 ± 0.23 action potentials (APs) to 4.4 ± 0.37 APs, F_1,9_=169.615, p<0.001; 60 pA pulse: from 3.2 ± 0.29 APs to 16.5 ± 0.80 APs, F_1,9_=214.846, p<0.001) (Figure 1 – figure supplement 1A-C). C-Fos immunohistochemistry also confirmed that intraperitoneal (i.p.) injection of CNO in 1mg/kg dose resulted in marked activation of BNST^vGAT^ neurons under baseline (homecage) condition (Figure 1 – figure supplement 1D: F_1,4_=145.377, p<0.001), which dose was able to induce behavioral effects, i.e. anxiogenic effect in the open field test (F_1,25_=5.255, p=0.030) as shown before (Mazzone *et al*, 2018), without affecting locomotor activity, i.e. no confounding effect on immobility (F_1,25_=0.559, p=0.461) (Figure 1 – figure supplement 1E). To test how BNST activation contributes to fear acquisition, we chemogenetically activated BNST^vGAT^ neurons during fear conditioning (Figure 1F; Figure – figure supplement 2). Elevated CS-induced fear recall (freezing) (F_1,15_=6.774, p=0.019) confirmed its functional role, although this effect diminished by the next extinction session 1 day later (i.e. Extinction recall: F_1,15_=1.275, p=0.276; Figure 1I). In contrast, contextual fear recall and freezing in a safe context during baseline period (i.e. before CS presentation as an index of contextual fear generalization) did not reveal any alterations (F_1,15_=0.025, p=0.875; F_1,15_=0.961, p=0.342, respectively; Figure 1H and I). Interestingly, chemogenetic activation did not affect freezing levels acutely during fear conditioning (Figure 1G: F_1,15_=0.041, p=0.841). Similarly, we tested how chemogenetic activation of BNST^vGAT^ neurons modulates CS-induced fear recall, or its generalization to safety cues using a differential auditory fear conditioning paradigm (Figure 1J). Experimental groups showed no difference in fear acquisition (F_1,19_=0.001, p=0.987) and could similarly differentiate between CS+ and CS-during a brief recall test (CS-/CS+: F_1,18_=0.451, p=0.510) (Figure 1K-L). In line with negative findings on c-Fos activity, we did not observe alteration in CS-induced freezing levels and CS+/CS-discrimination, when BNST^vGAT^ neurons were activated during fear recall (Figure 1M: CS+-induced recalls: F_1,19_=0.303, p=0.588; CS-/CS+: F_1,19_=0.043, p=0.837).

Taken together, BNST did not affect acute fear responses (exhibited either during conditioning or recall), supporting that predictable CS-induced fear expression is not dependent on BNST activity (Goode *et al*, 2019; Sullivan *et al*, 2004). However, it had significant impact on fear memory formation that *subsequently* manifested in enhanced fear levels. This finding initiated our next experiment to dissect the temporal dynamics (acquisition vs. consolidation), since chemogenetic activation has been reported to last for several hours (Roth, 2016).

### Chemogenetic activation of BNST facilitates fear learning via fear memory consolidation

Here, we replicated chemogenetic modulation of fear learning, but this time CNO was injected immediately after fear conditioning to activate BNST^vGAT^ neurons specifically during fear memory consolidation (Figure 2A-B; Figure 2 – figure supplement 1). This modulation replicated the enhancement of CS-induced fear recall (Figure 2E: F_1,15_=5.320, p=0.035), with no change in contextual recall (Figure 2D: F_1,17_=1.560, p=0.228). Since fear acquisition was similar between groups (Figure 2C: F_1,17_=0.311, p=0.584), enhanced BNST activity facilitated fear memory strength during consolidation, likely by modulating downstream regions of the fear circuitry.

**Figure 2.**
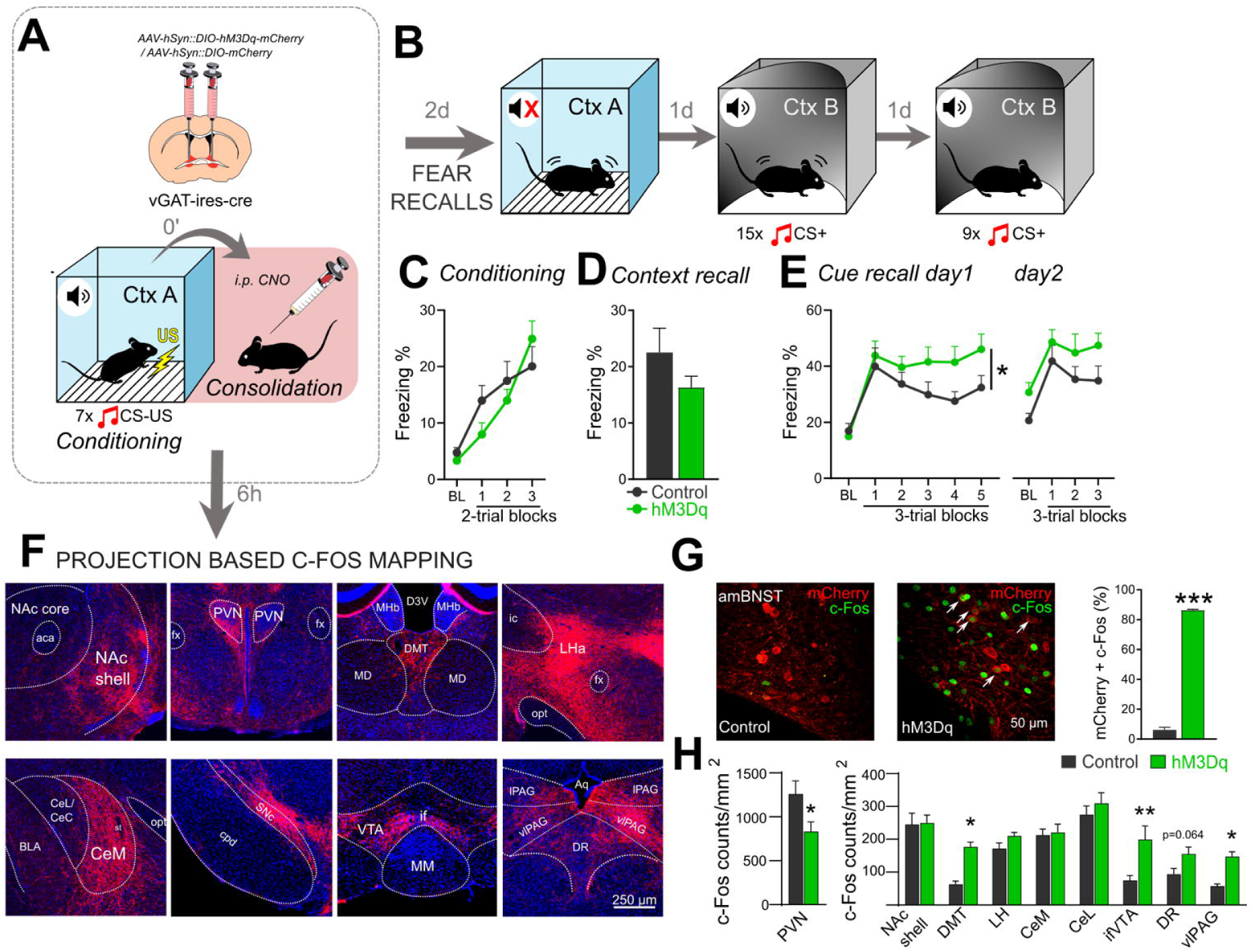
Chemogenetic activation of BNST^vGAT^ neurons during fear memory consolidation enhances CS-induced fear recall. Chemogenetic activation of BNST^vGAT^ neurons during fear memory consolidation resulted in enhanced cued fear recall, without altering contextual fear recall **(A-E).** (Control: n=10, hM3Dq: n=9). Representative wide-field fluorescence photomicrographs depicting major projection areas of BNST^vGAT^ neurons **(F)**, and assessed c-Fos expression 6 hours after chemogenetic activation (i.e. consolidation phase) **(G)**, where white arrows indicate activated hM3Dq-expressing BNST^vGAT^ neurons (mCherry+c-Fos). **(H)** C-Fos activity was significantly increased in the BNST, and DMT, vlPAG, ifVTA downstream regions, with additional decrease in PVN. *On freezing time curves each major tick depicts a 150 s block, starting with a pre- tone baseline period (BL). All data are represented as means ± s.e.m. Asterisks represent main effect of ANOVA: *p<0.05; **p<0.01, ***p<0.001. Abbreviations: aca-anterior commissure; amBNST: anteromedial BNST; Aq: <u>cerebral aqueduct;</u> BLA: basolateral amygdala; CeL/CeC: central amygdala, lateral/capsular part; CeM: central amygdala, medial part; cpd: cerebral peduncle; D3V: dorsal part of the third ventricle; DMT: dorsal midline thalamus; DR: dorsal raphe; fx: jornix; ic: internal capsule; LHA: Lateral hypothalamic area; MD: mediodorsal thalamus; MHb: medial habenula; MM: medial mammillary nucleus; Nae: nucleus accumbens; opt: optic tract; PVN: paraventricular nucleus of the hypothalamus, SNc: substantia nigra, pars compacta; st: stria terminalis; vlPAG/lPAG: periaqueductal gray, ventrolateral/lateral part; ifVTA: ventral tegmental area, interfascicular nucleus.*

To point out potential downstream targets mediating this effect, we mapped c-Fos activity 6 hours after fear conditioning when consolidation was chemogenetically enhanced by a similar design (Figure 2A and F). First, we confirmed marked activation of hM3Dq-expressing neurons in the BNST at 6 hours (86.36% vs. 6.09% in controls, Figure 2G; F_1,8_=1117.353, p<0.001). Second, we assessed c-Fos expression in densely innervated regions of the fear circuitry. We observed significant projections in nucleus accumbens (NAc) shell, dorsal midline thalamus (DMT), central amygdala (CeA, medial nucleus-CeM), lateral hypothalamus (LH), paraventricular nucleus of the hypothalamus (PVN), substantia nigra pars compacta (SNc), ventral tegmental area (VTA), dorsal raphe (DR), ventrolateral periaqueductal gray (vlPAG) (Figure 2F), in accordance with previous reports in rats and mice (Dong *et al*, 2001; Dong and Swanson, 2004, 2006a, b; Kodani *et al*, 2017). Among these regions, DMT, VTA (interfascicular part-ifVTA), and vlPAG exhibited increased c-Fos expression (F_1,13_=7.516, p=0.016; F_1,13_=8.191, p=0.013; F_1,14_=18.919, p<0.001, respectively), whereas PVN exhibited reduced expression levels (F_1,14_=5.208, p=0.038) in hM3Dq mice (Figure 2H; Figure 2 – figure supplement 2).

Although future studies targeting specific efferent pathways of BNST need to clarify how these specific downstream activities modulate fear learning, previous reports showed that these regions have significant impact on fear memory consolidation. Briefly, DMT is activated during aversive learning and exhibit plasticity-like changes in connection with prefrontal and amygdalar regions, particularly during fear memory consolidation (Do-Monte *et al*, 2015; Gao *et al*, 2020; Li *et al*, 2013; Penzo *et al*, 2014; Penzo *et al*, 2015). Amygdalar outputs initiate fear responses via vlPAG neurons (Dejean *et al*, 2015), which region has been recently shown to encode danger-cue associations and its probability (Ozawa *et al*, 2017; Wright *et al*, 2019a; Wright and McDannald, 2019b). Moreover, the avBNST-vlPAG pathway has been recently shown to modulate fear memory consolidation in the active avoidance paradigm (Lingg *et al*, 2020). Accordingly, BNST (as extension of CeA-DMT and CeA-vlPAG circuitry) could be a significant modulator of fear memory strength (Do-Monte *et al*, 2015; Ozawa *et al*, 2017; Penzo *et al*, 2015). Monoaminergic transmission (ifVTA) and glucocorticoids initiated from PVN can be also important elements how BNST exerts a tonic, plasticity-enhancing effect on fear memories as shown before (Dedic *et al*, 2018; Groessl *et al*, 2018; Lingg *et al*, 2020; Sengupta and Holmes, 2019).

In summary, our findings suggest that BNST modulates fear memory strength without direct impact on CS-induced fear response. Although this is in line with previous reports, directional contrasts with avBNST function, i.e. latter *reduced* subsequent fear response (Johnson *et al*, 2016; Johnson *et al*, 2019; Lingg *et al*, 2020), suggests different outcomes deriving from different BNST subregions and cell types. Indeed, marked heterogeneity and opposing impacts on hypothalamic-pituitary-adrenal (HPA) axis function and anxiety-like responses from different circuits of BNST have been documented (Choi *et al*, 2007; Crane *et al*, 2003; Gungor *et al*, 2016; Kim *et al*, 2013).

To interrogate the involvement of specific cell types of BNST in fear memory consolidation, we selectively modulated BNST^SOM^ and BNST^CRF^ neurons, which constitute two major GABAergic cell types in the BNST (Figure 3A-B) (Dedic *et al*, 2018; Nguyen *et al*, 2016; Ye and Veinante, 2019), and have been shown to drive passive and active forms of fear response in opposing manner in the CeA (Fadok *et al*, 2017). Moreover, CeA^SOM^ neurons enhance fear learning via connections with DMT and vlPAG, regions where we detected marked c-Fos hyperactivity (Li, 2019; Li *et al*, 2013; Penzo *et al*, 2014; Penzo *et al*, 2015). To test if CRF and SOM neurons mediate our memory consolidation enhancing effects in a differential manner, in the next step we applied SOM and CRF-specific chemogenetic activation and inhibition during the consolidation phase.

**Figure 3.**
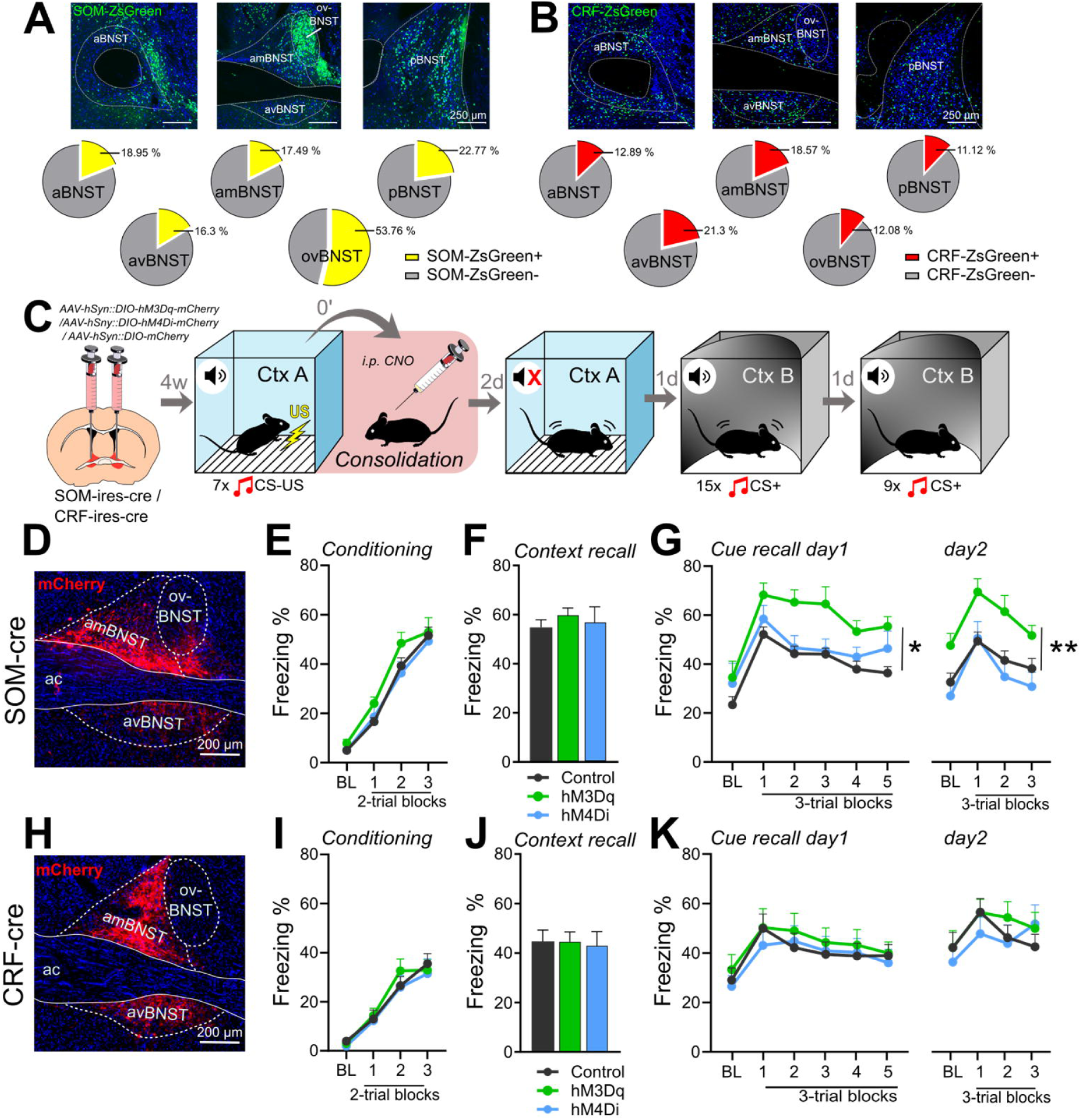
Chemogenetic activation of BNST^SOM^ neurons during fear memory consolidation enhances CS-induced fear recall. Distribution of SOM and CRF neurons in the BNST illustrated by representative single-plane confocal photomicrographs from reporter ZsGreen fluorescent protein expressing mouse lines (Gt(ROSA)26Sor-CAG/LSL-ZsGreen1), and their proportional quantification (for all neurons, NeuN+) across subregions **(A-B).** Experimental design for chemogenetic modulation of BNST^SOM^and BNST^CRF^ neurons during fear memory consolidation **(C).** Chemogenetic activation of BNST^SOM^ neurons during fear memory consolidation resulted in enhanced cued fear recall, without altering contextual fear recall **(D-G, green)** (Control: n=8, hM3Dq: n=7). In contrast, chemogenetic inhibition of BNST^SOM^ neurons **(D-G, blue**, hM4Di: n=9), or chemogenetic modulations of BNST^CRF^ neurons had no impact on fear recalls **(H-K)** (Control: n= 15, hM3Dq: n= 12, hM4Di: n= 13). *On freezing time curves each major tick depicts a 150 s block, starting with a pre-tone baseline period (BL). All data are represented as means ± s.e.m. Asterisks represent main effect of ANOVA: *p<0.05; **p<0.01. Abbreviations: ac: anterior commissure, amBNST: anteromedial BNST, avBNST: anteroventral BNST, ovBNST: oval nucleus of the BNST; pBNST: posterior BNST.*

### BNST^SOM^, but not BNST^CRF^, neurons facilitate fear memory consolidation

Chemogenetic activation of BNST^SOM^ neurons in Som-ires-cre mice during fear consolidation (Figure 3C and D, Figure 3 – supplement 2) replicated our BNST^vGAT^ findings by resulting in enhanced fear recall which was even more pronounced and persistent (Figure 3G, Cue recall day1: F_2,30_=8.067, p=0.001, Tukey post hoc p=0.001 and p=0.391 control vs. hM3Dq and hM4Di, respectively; day2/Extinction recall: F_2,29_=5.547, p=0.009, Tukey post hoc p=0.021 and p=0.773 control vs. hM3Dq and hM4Di, respectively; CtxB BL’s for day1: F_2,31_=0.769, p=0.471 and day2: F_2,29_=4.225, p=0.024). Again, contextual fear recall was not changed (Figure 3F, F_2,31_=0.160, p=0.852), and fear conditioning was similar between groups (Figure 3E, F_2,29_=1,512, p=0.237). Interestingly, chemogenetic inhibition did not affect any forms of fear recall, despite its potential to lower neuronal activity indicated by c-Fos (F_2,10_=93,361, p<0.001, Tukey post hoc p=0.008; Figure 3 – figure supplement 1). In contrast to BNST^SOM^ neurons, chemogenetic activation (and inhibition) of BNST^CRF^ neurons during fear consolidation in crh-ires-cre mice (Fear conditioning: F_2,37_=0.205, p=0.815; Figure 3C; Figure 3 – figure supplement 2) had no effect on contextual (Figure 3J, F_2,37_=0.041, p=0.959), and CS-induced fear recalls (Figure 3K, Cue recalls: F<0.270, p>0.765), or fear generalization (CtxB BL’s: F<0.416, p>0.662).

Latter findings also suggest that SOM positive neurons in the central extended amygdala have a rather uniform impact on fear-like responses, shifting it towards passive forms (Fadok *et al*, 2017; Li *et al*, 2013; Yu *et al*, 2016), although BNST^SOM^ neurons may not be inevitable for intact fear learning as chemogenetic inhibition showed no effect. However, BNST repeatedly exerted a modulatory role on the memory formation of the US-CS association, which may be relevant in vulnerable populations exhibiting excessive fear learning. Noteworthy, a similar one-directional effect was observed following optogenetic manipulation of specific pathways originating from BNST (Lingg *et al*, 2020). Negative findings on BNST^CRF^ activation confirm previous reports (Marcinkiewcz *et al*, 2016), however, we cannot exclude that our paradigm was not sufficient to reveal ‘active-defensive’ phenotype regulated by CRF, i.e. escape behavior, which was hardly present during conditioning or recall (<1%, sporadic occurrence), and likely requires specific stimuli and settings to manifest as shown before (Fadok *et al*, 2017). Although CRF signaling in BNST has been reported as anxiogenic, its impact seems less prominent in fear learning and may originate from CeA inputs (Asok *et al*, 2018; Pomrenze *et al*, 2019; Regev *et al*, 2011).

Considering downstream target areas of BNST^SOM^ and BNST^CRF^ neurons (and BNST^vGAT^), we could observe minimal differences between these cell types (Figure 3 – figure supplement 3 and Figure 2F), which was also highly overlapping with reported CeA projections (Hartley *et al*, 2019; Li, 2019), and corresponded with previous descriptions of BNST^CRF^ projections (Dabrowska *et al*, 2016; Dedic *et al*, 2018). Different impact of SOM and CRF may originate from different contacts with specific cell types, which is to be clarified by future studies. Considering their intra-BNST distribution, we observed moderate differences between CRF and SOM expression in medial regions (∼15-20%; Figure 3A-B), however, SOM neurons showed significant dominance in the oval nucleus and posterior region indicated by cre-dependent reporter Zsgreen fluorescent protein expression in crossed mouse lines (Gt(ROSA)26Sor-CAG/LSL-ZsGreen1). Noteworthy, we detected very limited expression of DREADDs in the oval nucleus in our experiments.

## Conclusion

Taken together, our study points out a specific role of BNST in the regulation of fear learning. Namely, BNST^SOM^ neurons facilitate CS-US fear memory formation and its strength (likely via plastic changes in downstream regions) without affecting acute fear reactivity and expression. These data help to better understand how amygdala and BNST exert complementing functions in the fear circuitry, which may resolve some controversies between early and recent models explaining functional division across extended amygdala regions (Davis *et al*, 2010; Fox and Shackman, 2019; Goode *et al*, 2019; Gungor *et al*, 2016). Finally, BNST hyperactivity (as modeled in the present study) may translate into individual vulnerability in humans (i.e. enhanced activity of BNST in individuals with anxiety disorders or high anxiety traits; (Avery *et al*, 2016; Brinkmann *et al*, 2018)).

## Materials and Methods

### Subjects

Adult (>8 weeks old) male mice from the following strains were used in the present study: C57Bl/6J, vGAT-ires-cre, CRH-ires-cre and SOM-ires-cre mice (all strains from Jackson Laboratory, USA) (Taniguchi *et al*, 2011; Vong *et al*, 2011). To visualize CRF and SOM neurons (by reporter fluorescent proteins) in the BNST, we crossed CRH-ires-cre and SOM-ires-cre mice with Gt(ROSA)26Sor-CAG/LSL-ZsGreen1 mice (Jackson Laboratory, USA). All animals were group-housed (3-4 mice/cage) in Plexiglass chambers at constant temperature (22 ± 1 °C) and humidity (40–60%), under a reverse circadian light-dark cycle (lights-off at 7:00 a.m., lights-on at 7:00 p.m.). Behavioral experiments were performed during the first half of the active (dark) cycle. Mice were isolated 3 days before fear conditioning and kept single-housed during testing to prevent social buffering/modulatory effects. Regular laboratory chow (Sniff, Soest, Germany) and water were available *ad libitum*.

### Stereotaxic surgery for viral gene transfer

Mice underwent stereotaxic surgery to bilaterally inject virus constructs into the BNST (anterior-posterior +0.8 mm, medio-lateral ±0.8 mm, dorso-ventral –4.2 mm to Bregma; (Paxinos and Franklin, 2001)). Animals were anesthetized with a ketamine-xylazine solution (16.6 mg/ml ketamine and 0.6 mg/ml xylazine-hydrochloride in 0.9% saline, 10 ml/kg body weight i.p.) and placed in a stereotaxic frame (David Kopf Instruments, Tujunga, CA, USA). Viral vectors (20-40 nl volume/hemisphere) were microinjected through a glass pipette (tip diameter: 20–30 μm) at a rate of 100 nl/minute by using a Nanoject II precision microinjector pump (Drummond, Broomall, PA, USA). The pipette was left in place for an additional 5 min to ensure diffusion before slow retraction. After the surgeries, mice received buprenorphine injection (Bupaq; 0.1 mg/kg) subcutaneously as analgesic treatment. Behavioral experiments were conducted 4-6 weeks after virus injection to allow time for DREADD expression.

### Virus vectors

Adeno-associated viruses (AAVs) carrying Cre-inducible (double-inverse orientation; DIO) transgenes were purchased from Addgene (Watertown, MA, USA). We used stimulatory AAV8-hSyn::DIO-hM3Dq-mCherry (4.0e12 GC/ml titer, #44361), inhibitory AAV8-hSyn::DIO-hM4Di-mCherry (1.9e13 GC/ml titer, #44362) DREADD constructs, and inactive control fluorophore AAV8-hSyn::DIO-mCherry (4.1e12 GC/ml titer, #50459).

### Drugs

Designer receptor-ligand clozapine-N-oxide (CNO, Tocris Bioscience; 4936, CAS No: 34233-69-7) was dissolved in 0.9 % saline solution at a concentration of 0.1 mg/ml and administered i.p. at a dose of 1 mg/kg 40 min before testing (in case of open field, fear acquisition or fear recall modulation), or immediately after fear conditioning (in case of modulation of fear memory consolidation).

### Behavioral testing

#### Open field test

Animals were tested in the open field arena to verify the behavioral, i.e. anxiogenic effect of chemogenetic stimulation of BNST. The open-field arena was made of white plastic (40 × 30 × 15 cm), cleaned with water between animals. Mice were placed in the corner and allowed to explore the arena for 10 minutes. The inner 20 × 15 cm zone was considered as center, and time spent here was an index of anxiety. Total distance moved was the index of locomotor activity. Behavior was measured using EthoVision XT 13 software (Noldus Information Technology, Wageningen, Netherlands). Mice underwent two open field tests 1 week apart: first one measuring baseline open field activity (without CNO treatment), second one measuring open field activity with chemogenetic activation (1 mg/kg i.p. CNO treatment 40 min before testing). Control mice expressing control fluorophore received the same CNO treatment before testing. Latter applies for all tests of our studies.

#### Auditory fear conditioning and recall testing

Auditory fear conditioning was started 4-6 weeks after viral surgeries (for visualized schematics, see Fig.1F). On day 1, mice were placed into a clear plexiglass chamber (25 × 25 × 30 cm) with electrical grid floor (Coulbourn Instruments, Holliston, MA, USA) used to deliver footshocks. The chamber was cleaned with 20% ethanol between animals. Fear conditioning was performed with maximum light intensity in the test room (700 lux). This setting was considered as the conditioning context (CtxA: light on, ethanol odor, specific box with grid floors). Following a 150 s baseline period, mice were presented with seven auditory stimuli as conditioned stimuli (CS+: 80 dB pure tones at 7 kHz, SuperTech Instruments, Pecs, Hungary) of 30 s duration spaced with pseudorandom interstimulus intervals (ISIs, ranged between 60–90 s). All CS+ co-terminated with a 1 s scrambled footshock as unconditioned stimulus (US: 0.7 mA).

On day 3, mice were exposed briefly to context A for five minutes without CS+ to study contextual fear recall without inducing fear extinction. On days 4 and 5, mice were subjected to sessions of auditory fear recall in an altered context B (CtxB). Context B was altered in all dimensions to differ from Context A (i.e. experimenter, room, 20 lux red light, plastic floor instead of grids, plastic inserts changing the shape of the chamber, cleaning with soapy water with fruit odor). Both sessions started with a 150 s baseline period (no CS+ presented) to measure fear response in a safe context (conditioned vs. safe context fear responses as an index of generalization). Baseline period was followed by 15 or 9 CS+ (30 s duration, spaced with 30 s ISIs) on days 4 and 5, respectively, to measure cue-dependent fear recall, as well as within- and between-session fear extinction (Figure 1F). Time spent with freezing was considered as an index of fear, which was analyzed using EthoVision XT 13 software. Software parameters and thresholds were set and optimized to reach R > 0.90 correlation with handscoring by an experimenter blind to treatments.

#### Differential auditory fear conditioning and recall testing

Fear conditioning and recall testing took place in CtxA and B, respectively (Figure 1J). In this paradigm, we presented two types of auditory stimuli with 30 s duration, i.e. 7 kHz pure tone and white noise pips. They were used as conditioned and unconditioned stimuli (CS+ and CS-, randomized between animals, and counterbalanced between groups). On day 1, mice were habituated to auditory cues in CtxA, i.e. after a 150 s baseline period 4 of each cue were delivered in alternating order with 30 s ISIs (not shown in Figure 1). Next day, fear conditioning was performed in CtxA: after 150 s baseline period, 7 CS+ and 7 CS- (30 s duration, 30s ISIs) were presented in alternating order. All CS+, but none of CS-, were co-terminated with 1 s scrambled footshock (0.5 mA). We used lower shock intensity to achieve better discriminative learning in mice based on previous studies (Duvarci *et al*, 2009; Kim and Cho, 2017; Sanford *et al*, 2017). Two days later cue-dependent fear recall and discrimination between CS+ and CS- was briefly tested by exposing subjects to 4 CS+ and 4 CS- in alternating order in CtxB (30s duration, 30s ISIs). Next day, cue-dependent fear recall (with CS+/CS-discrimination) and fear extinction were tested following chemogenetic activation of BNST (1 mg/kg i.p. CNO; 40 min pre-injection time) by exposing subjects to 15 CS+ and 4 CS- in alternating order in CtxB (after 150 s baseline period). We also tested extinction recall by exposing mice to 9 CS+ in CtxB 24 hrs later. Analysis of freezing behavior was carried out as above.

### Ex vivo electrophysiology and slice preparation

For validation of hM3Dq mediated depolarization, current-clamp recordings were performed on brain slices obtained from vGAT-cre mice (postnatal days 90-110, matching experimental age) expressing hM3Dq or control fluorophore in BNST (n=4/group; Figure 1 – figure supplement 1A-C). Animals were given 4-6 weeks to express transgenes after virus injection. After decapitation, the brain was removed rapidly and immersed in ice-cold low-sodium solution (in mM: saccharose 205.0, KCl 2.5, NaHCO_3_ 26.0, CaCl_2_ 1.0, MgCl_2_ 5.0, NaH_2_PO_4_ 1.25, and glucose 10) bubbled with a mixture of 95% O_2_ and 5% CO_2_. Coronal sections of the BNST were sliced at 250 μm on a VT-1000S Vibratome (Leica Microsystems, Wetzlar, Germany) in the low-sodium solution. The slices were transferred into artificial CSF (aCSF; in mM: NaCl 130.0, KCl 3.5, NaHCO_3_ 26.0, CaCl_2_ 2.5, MgSO_4_ 1.2, NaH_2_PO_4_ 1.25, and glucose 10) saturated with O_2_/CO_2_, and kept in it for 1 h to equilibrate. The initial temperature of aCSF was 33°C, which was left to cool to room temperature during equilibration. Electrophysiological recording, during which the brain slices were oxygenated by bubbling the aCSF with O_2_/CO_2_ gas, was performed at 33°C. An Axopatch 200B patch-clamp amplifier, a Digidata-1322A Data Acquisition System, and pCLAMP version 10.4 software (Molecular Devices, USA) were used for recording.

Recordings were performed under visual guidance using a BX51WI IR-DIC microscope (Olympus, Japan) located on a S’Table antivibration table (Supertech Instruments, Pecs, Hungary) to detect mCherry-positive cells.

The patch electrodes (outer diameter, 1.5 mm, thin wall; Hilgenberg) were pulled with a Flaming-Brown P-97 puller (Sutter Instrument, Novato, CA, USA) and polished with an MF-830 Microforge (Narishige, Tokyo, Japan). The pipette solution contained the following (in mM): K-gluconate 130, NaCl 10, KCl 10, MgCl_2_ 0.1, HEPES 10, EGTA 1, Mg-ATP 4, and Na-GTP 0.3 (pH 7.3 with KOH). Osmolarity was adjusted to 295–300 mOsm with sorbitol. Neurons were recorded in whole-cell mode, the intrapipette solution contained 0.2% biocytin. To record CNO-induced depolarization of hM3Dq expressing neurons, 10 μM CNO was bath applied for 10 min after a 3-min baseline in the presence of 660 nM TTX. Firing was induced by 30 or 60 pA current pulses (900 ms), respectively, 5 min after onset of CNO application. Similar recordings were performed on control mice (expressing inactive control fluorophore DREADD). Recordings were stored and analyzed off-line using the Clampfit module of the PClamp version 10.4 software (Molecular Devices, San Jose, CA, USA).

### Immunohistochemistry and image analysis

#### Tissue processing

Mice were anesthetized with a ketamine-xylazine solution (16.6 mg/ml and 0.6 mg/ml, respectively) and transcardially perfused with ice-cold phosphate-buffered saline (PBS), followed by ice-cold paraformaldehyde (PFA; 4% in PBS). Brains were rapidly removed and postfixed in 4% PFA at 4 °C, and cryoprotected in 30% sucrose solution in PBS before slicing. 30 μm coronal sections were collected on a sliding microtome and stored in a cryoprotectant solution (containing 20% glycerin, 30% ethylene glycol) at −20 °C until immunohistochemical analysis.

#### Verification of virus extensions

We labeled mCherry by immunohistochemistry using primary antibody against red fluorescent protein (RFP) to verify virus expression in the BNST (Figure 1 – figure supplement 1D). Briefly, after several rinses in PBS, sections (90 μm apart) were incubated in PBS containing 0.3% Triton X-100 (TxT, Sigma-Aldrich) and 0.3% H_2_O_2_ for 30 min followed by 2% bovine serum albumin (BSA, Sigma-Aldrich) diluted in PBS for 1 hour. Primary antibody solution (1:4000 rabbit anti-RFP, #600-401-379, Rockland, Limerick, PA, USA; diluted in PBS containing 2% BSA and 0.1% Triton-X) was left over on the slices for 2 days at 4°C. After several rinsing with PBS, slices were incubated in biotin-conjugated donkey anti-rabbit secondary antibody (1:1000 in 2% BSA and PBS, #711-065-152, Jackson ImmunoResearch, Cambridgeshire, United Kingdom) for 2 hours. Labeling was amplified by avidin–biotin complex (1:1000; Vector Laboratories, Burlingame, CA, USA) by incubation for 1 h at room temperature. The peroxidase reaction was developed in the presence of diaminobenzidine tetrahydrochloride (0.2 mg/ml), nickel–ammonium sulfate (0.1%), and hydrogen peroxide (0.003%) dissolved in Tris buffer. Sections were mounted onto gelatin-coated slides, dehydrated, and coverslipped with DPX Mountant (Sigma-Aldrich/Merck, Darmstadt, Germany). Regions of interest were digitalized by an Olympus DP70 Light Microscope and CCD camera system. All animals with virus extension outside of the BNST were excluded from the analysis. Generally, mCherry-positive cell bodies were observed along the whole rostro-caudal axis of the BNST, including anteroventral, anteromedial, and posterior regions of the BNST, with limited expression in the oval nucleus (Figure 1-3 – figure supplements).

#### Viral tracing

We also used anti-RFP fluorescent immunolabeling and confocal microscopy to assess projections of BNST^CRF^, BNST^SOM^, and BNST^vGAT^ neurons. The protocol was slightly modified as above: non-specific binding sites were blocked by 10% normal goat serum (NGS diluted in TBS, Jackson ImmunoResearch) for 1 hour, and slices were incubated in the primary antibody solution for 3 days (monoclonal rabbit anti-RFP IgG 1:1000, #600-401-379, Rockland; 0,15% TxT in TBS). Several rinsing with TBS was followed by incubation in secondary antibody solution for 2 hours (1:500 Cy3 conjugated goat anti-rabbit IgG, #134845, Jackson ImmunoResearch).

#### C-Fos immunohistochemistry

We used c-Fos immunohistochemistry to assess neuronal activity in BNST and its downstream regions at different time points of fear learning. Mice were anesthetized (with ketamine-xylazine mixture) and transcardially perfused 90 min after either testing or CNO injection (in case of homecage condition to verify chemogenetic activation of BNST in vivo) as described above. We used fluorescent immunolabeling against c-Fos and RFP as described above (1:2000 guinea-pig polyclonal anti-c-Fos IgG, #226004, Synaptic Systems with monoclonal rabbit anti-RFP IgG 1:1000, #600-401-379, Rockland), which were detected by fluorescent-conjugated antibodies (1:500 Cy3 conjugated donkey anti-rabbit, #134845, Jackson ImmunoResearch, and 1:500 Alexa-488 conjugated donkey anti-guinea-pig, #S32354, ThermoFisher Scientific, Waltham, MA, USA).

#### Biocytin labeling

Biocytin labeled neurons following electrophysiological recording were confirmed to be mCherry positive by consequent immunolabeling and confocal microscopic analysis. Slices were incubated in 4% PFA in 0.1 M phosphate buffer (PB; pH 7.4) overnight. Next day, after PBS rinses, sections were incubated with 10% NGS and 0.3% TxT in PBS for 1 hour, followed by primary antibody solution incubation overnight (1:2000 monoclonal rabbit anti-RFP IgG, #600-401-379, Rockland, in 3% NGS and 0.3% TxT). After PBS rinses, primary antibodies were detected by Alexa 488-conjugated streptavidin (1:500 Alexa-488 conjugated donkey anti-guinea-pig, #S32354, ThermoFisher Scientific) and Alexa 594-conjugated goat anti-rabbit IgG (1:200, #A11012, ThermoFisher Scientific) dissolved in PBS containing 3% NGS and 0.3% TxT, incubated overnight. After several rinsing in PBS, sections were mounted on glass slides and coverslipped using Mowiol4 –88 (Sigma-Aldrich/Merck).

#### SOM- and CRF-Zsgreen assessment

To quantify the distribution of SOM and CRF neurons in BNST subregions, neuronal somata were visualized by NeuN immunolabeling (1:1000, monoclonal mouse anti-NeuN IgG, #MAB377, /Merck, Darmstadt, Germany) in CRH-ires-cre: and SOM-ires-cre::Gt(ROSA)26Sor-CAG/LSL-ZsGreen1 crossed mouse lines. Primary antibody was incubated overnight in 2% normal donkey serum and 0.1% TxT). Next day, after PBS rinses, primary antibody was detected by Cy3-conjugated donkey anti-mouse IgG (1:1000, # 715-165-151 Jackson ImmunoResearch). After several rinsing in PBS, sections were mounted on glass slides and coverslipped using Mowiol4–88 (Sigma-Aldrich/Merck).

#### Microscopy and quantification of labeling

All imaging and quantification were performed by experimenters blind to treatments. As mentioned above Ni-DAB-stained sections were digitalized by an Olympus DP70 Light Microscope and CCD camera system. Fluorescent c-Fos labeling was imaged using C2 Confocal Laser-Scanning Microscope (Nikon CFI Plan Apo VC60X/NA 1.40 Oil objective, z step size: 0.13 μm, xy: 0.08 μm/pixel and CFI Plan Apo VC20X/N.A. 0.75, xy: 0.62 μm/pixel, Nikon Europe, Amsterdam, The Netherlands), whereas projections of BNST (anti-RFP) were imaged using a Panoramic Digital Slide Scanner (Zeiss, Plan-Apochromat 10X/NA 0.45, xy: 0.65 μm/pixel, Pannoramic MIDI II; 3DHISTECH, Budapest, Hungary) equipped with LED (Lumencor, SPECTRA X light engine). To assess c-Fos counts and ZsGreen positive BNST^SOM^/BNST^CRF^ neurons, we delineated regions of interest or used fixed rectangular/oval frames on fluorescent pictures using CaseViewer 2.3 software (3DHISTECH, Budapest, Hungary). All c-Fos results are presented as c-Fos counts normalized for mm^2^, which were counted manually using standardized settings (contrast, intensity) across subjects and regions.

### Statistics

Data are expressed as mean ± standard error of the mean. Differences between groups were evaluated by one-way or repeated-measure ANOVA (time/block as within subject factor), followed by Tukey’s post hoc analyses using Statistica software (Tibco, Palo Alto, CA, USA). Ex vivo electrophysiological data were also analyzed by Student’s one-sample t-test. Significance level was set at p<0.05 throughout, however, all p values are indicated with exact numbers.

## Acknowledgments

This study was supported by National Research, Development and Innovation Office Grants (Hungary) #FK129296 and #PD116589 (for MT), and the Hungarian Brain Research Program grant #2017-1.2.1-NKP-2017-00002 (for EM). This work was also supported by New National Excellence Program Grants #UNKP-18-3-III-SE-22, #UNKP-19-2-I-ELTE-574 (for LB and HS). We thank all the core facilities of our institute for their supportive help: the Behavioral Studies Unit for help with behavioral testing (Dr. Kornél Demeter), the Nikon Microscopy Center for help with microscopy (Dr. László Barna and Dr. Csaba Pongor), the Medical Gene Technology Unit for help with mouse lines. We also thank for Beata Barsvari and Nikolett Bakos for technical assistance.

## Additional information

### Funding

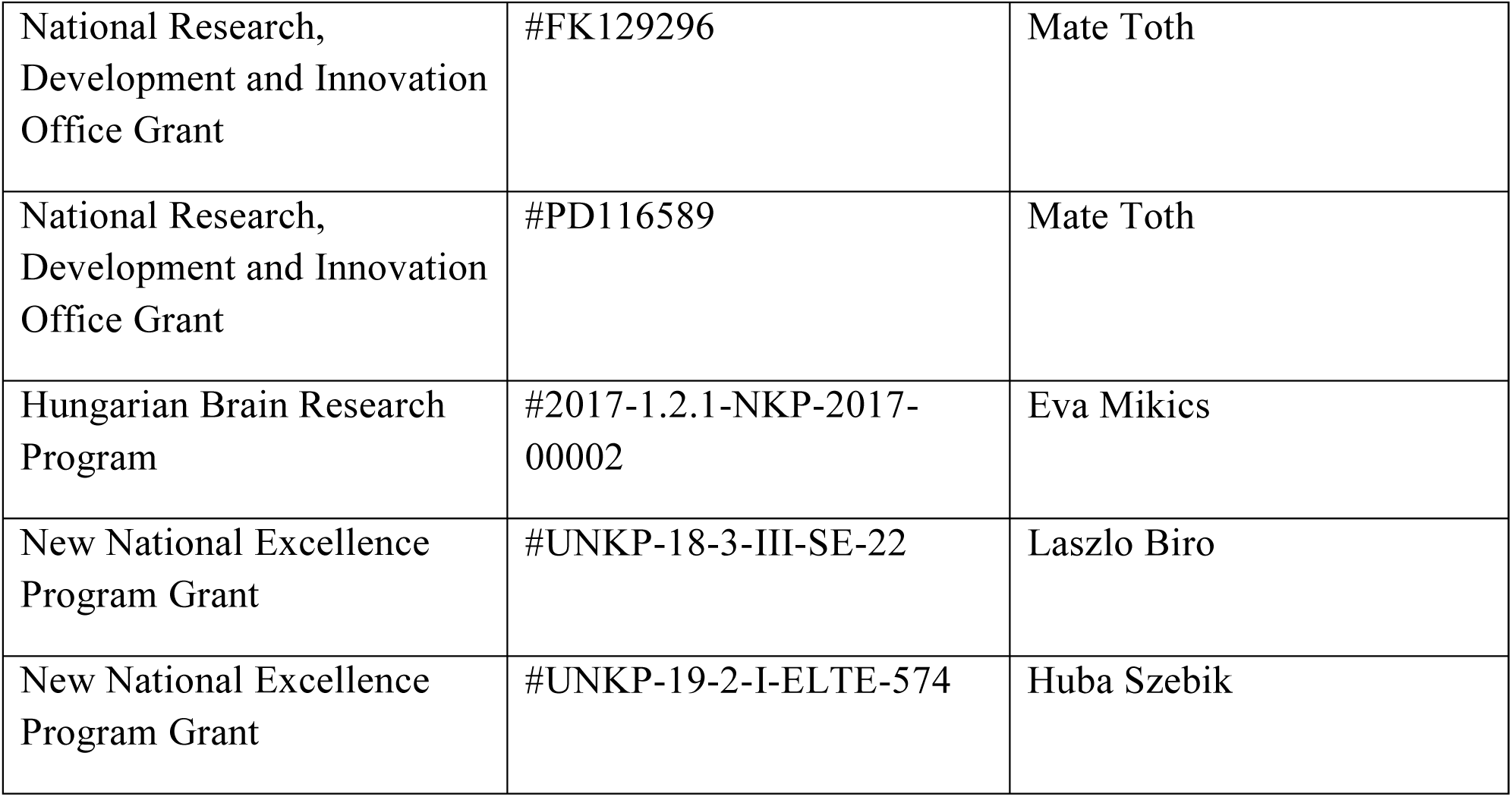

The funders had no role in study design, data collection and interpretation, or the decision to submit the work for publication.

### Author contributions

Biborka Bruzsik, Laszlo Biro, Conceptualization, Investigation, Methodology, Data analysis, Visualization, Writing – original draft;

Dora Zelena, Eszter Sipos, Huba Szebik, Klara Rebeka Sarosdi, Orsolya Horvath, Cintia Klaudia Finszter, Investigation;

Imre Farkas, Veronika Csillag, Investigation, Data analysis;

Eva Mikics, Supervision, Resources, Funding acquisition Mate Toth, Conceptualization, Supervision, Data analysis, Funding acquisition, Writing – original draft

### Ethics

All procedures were conducted in accordance with the guidelines of European Communities Council Directive of 2010 (2010/63/EU) and were reviewed and approved by the Animal Welfare Committee of the Institute of Experimental Medicine, Hungary.

---

**Figure 1 – figure supplement 1.**
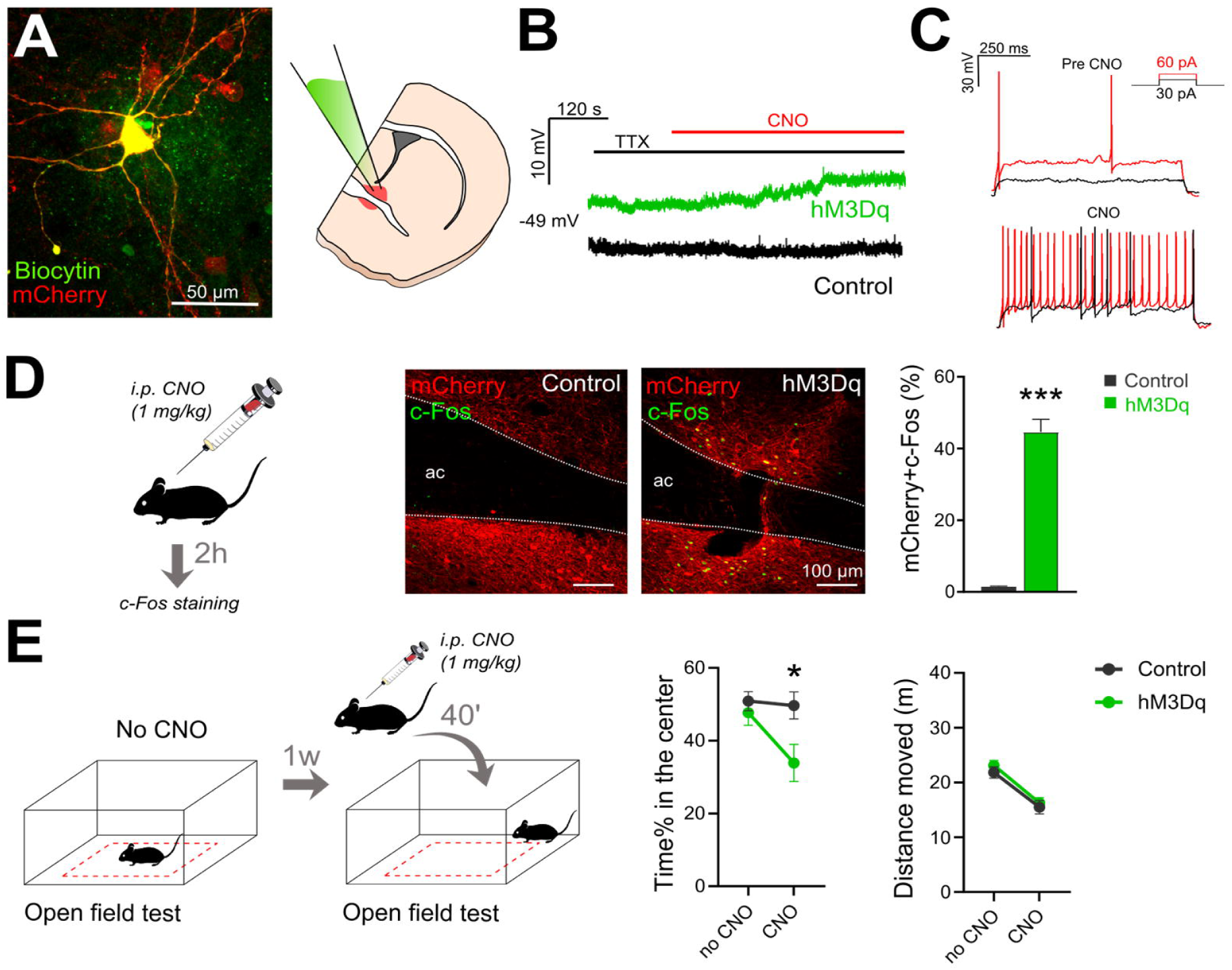
*In vitro* and *in vivo* verification of hM3Dq-mediated activation of BNST^vGAT^ neurons. Schematics of BNST slice recordings and representative photomicrograph of a biocytin-filled BNST^vGAT^ neuron expressing hM3Dq **(A)**, with a representative trace of hM3Dq mediated depolarization in the presence of TTX and CNO, which was absent in BNST^vGAT^ neurons expressing control fluorophore **(B).** CNO administration also elevated the frequency of action potentials evoked by depolarizing current steps and decreased the rheobase **(C).** Intraperitoneal injection of CNO (I mg/kg) under baseline (homecage) condition induced significant c-Fos expression in hM3Dq-mCherry-expressing neurons (Control: n=3, hM3Dq: n=3 mice) **(D).** CNO (I mg/kg) increased anxiety in the open field test without locomotor effects **(E).** (Control: n=l5, hM3Dq: n=l2 mice) *All data are represented as means* ***±*** *s.e.m. Asterisks represent main effect of ANOVA: *p<0.05; ***p<0.001.*

**Figure 1 – figure supplement 2.**
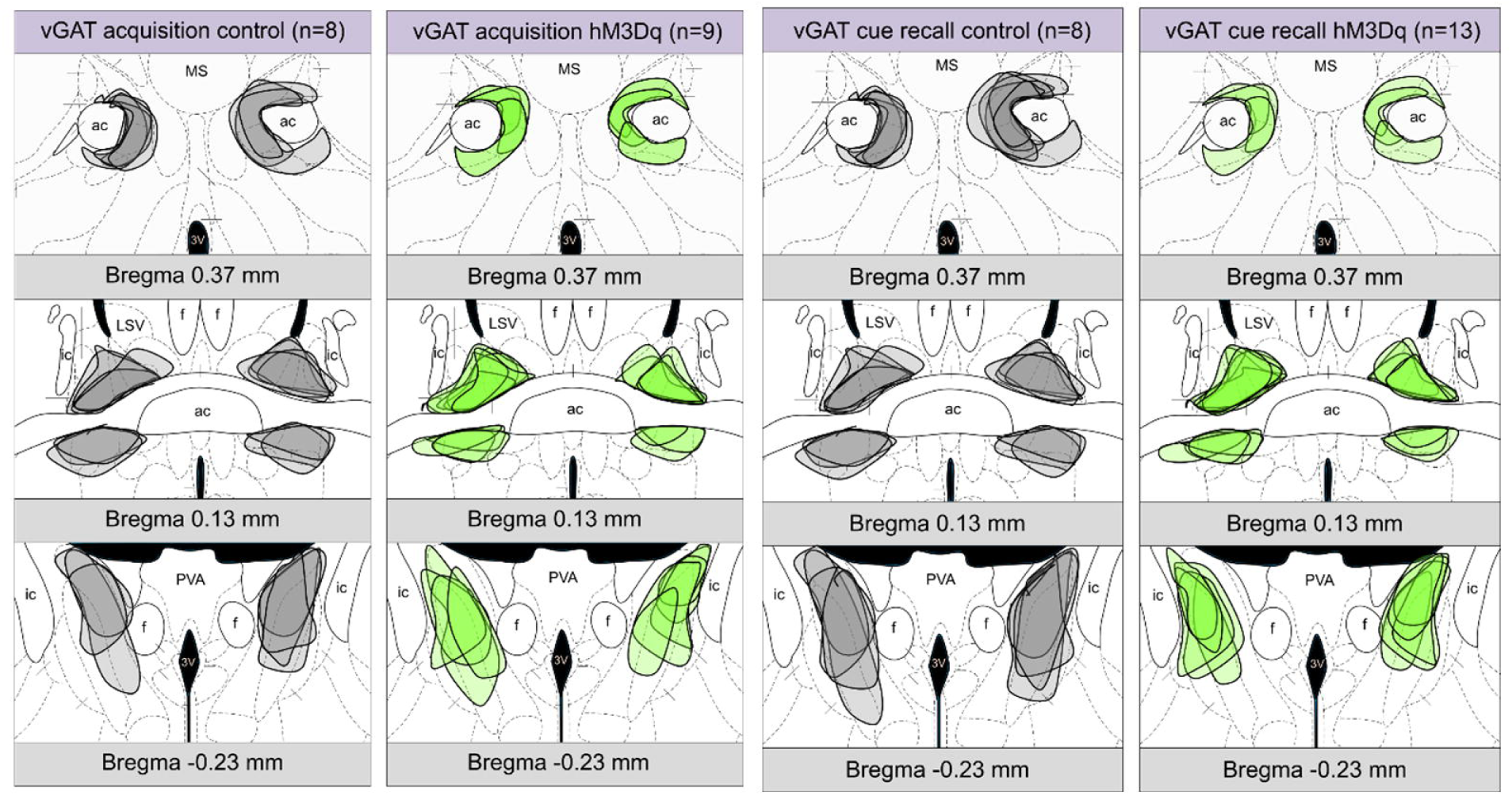
Illustration of AAV-associated mCherry expression in the BNST from control (gray shaded) and hM3Dq-mCherry expressing vGAT-ires-cre mice, which were CNO-injected during fear conditioning and cued fear recall. Diagrams were modified from the Allen Mouse Brain Atlas, Allen Institute for Brain Science (http://mouse.brainmap.org/). *Abbreviations: 3V: third ventricle, ac: anterior commissure, f:fornix, ic: internal capsule, LSV-lateral septal nucleus, ventral part, MS: medial septum, PVA: paraventricular nucleus of thalamus.*

**Figure 2 – figure supplement 1.**
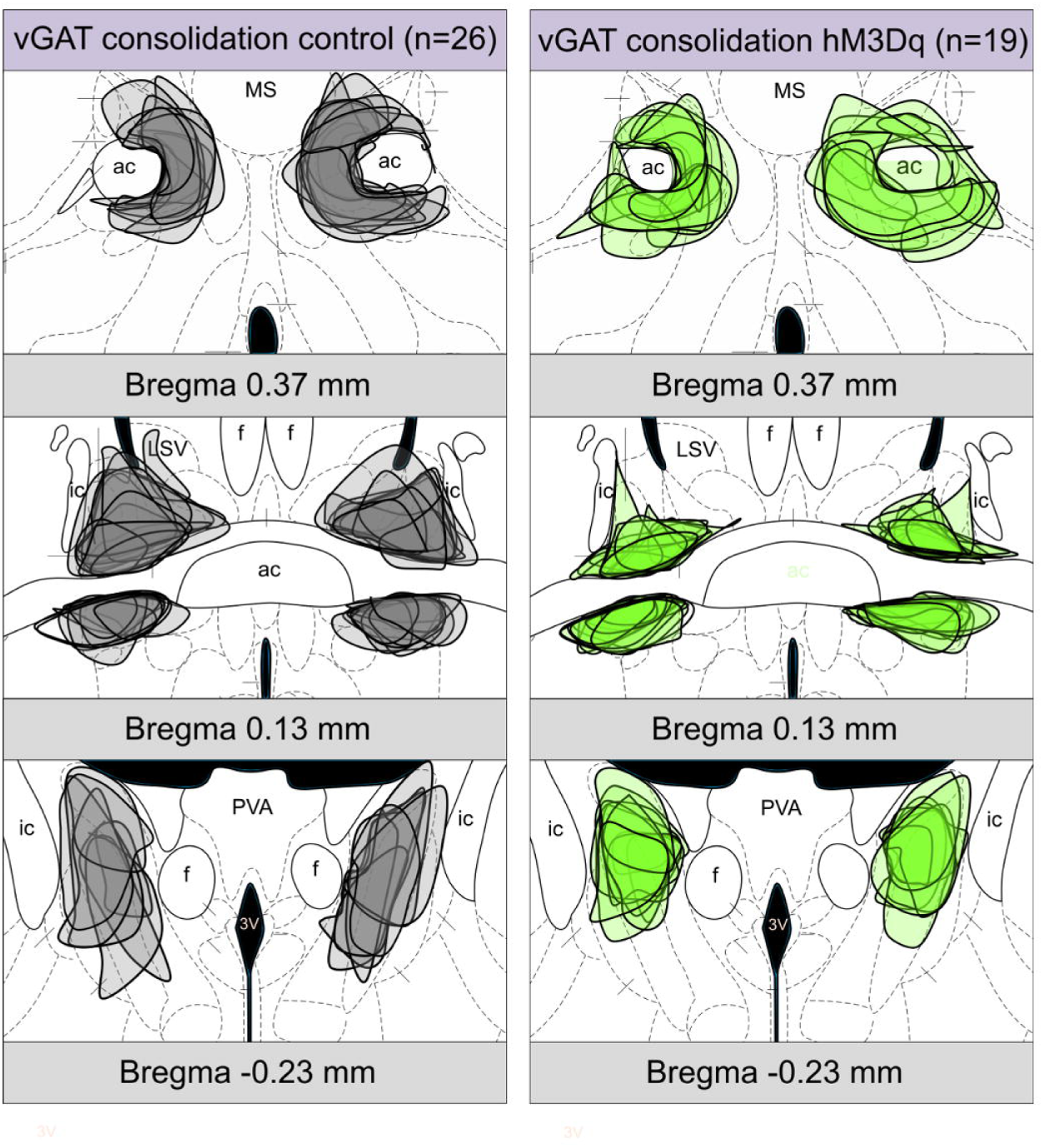
Illustration of AAV-associated mCherry expression in the BNST from control (gray shaded) and hM3Dq-mCherry expressing vGAT-ires-cre mice, which were CNO-injected during the consolidation phase. Diagrams were modified from the Allen Mouse Brain Atlas, Allen lnstitute for Brain Science (http://mouse.brainmap.org/). *Abbreviations: 3V: third ventricle, ac: anterior commissure, f: fornix, ic: internal capsule, LSV: lateral septal nucleus, ventral part, MS: medial septum, PVA: paraventricular nucleus of thalamus.*

**Figure 2 – figure supplement 2.**
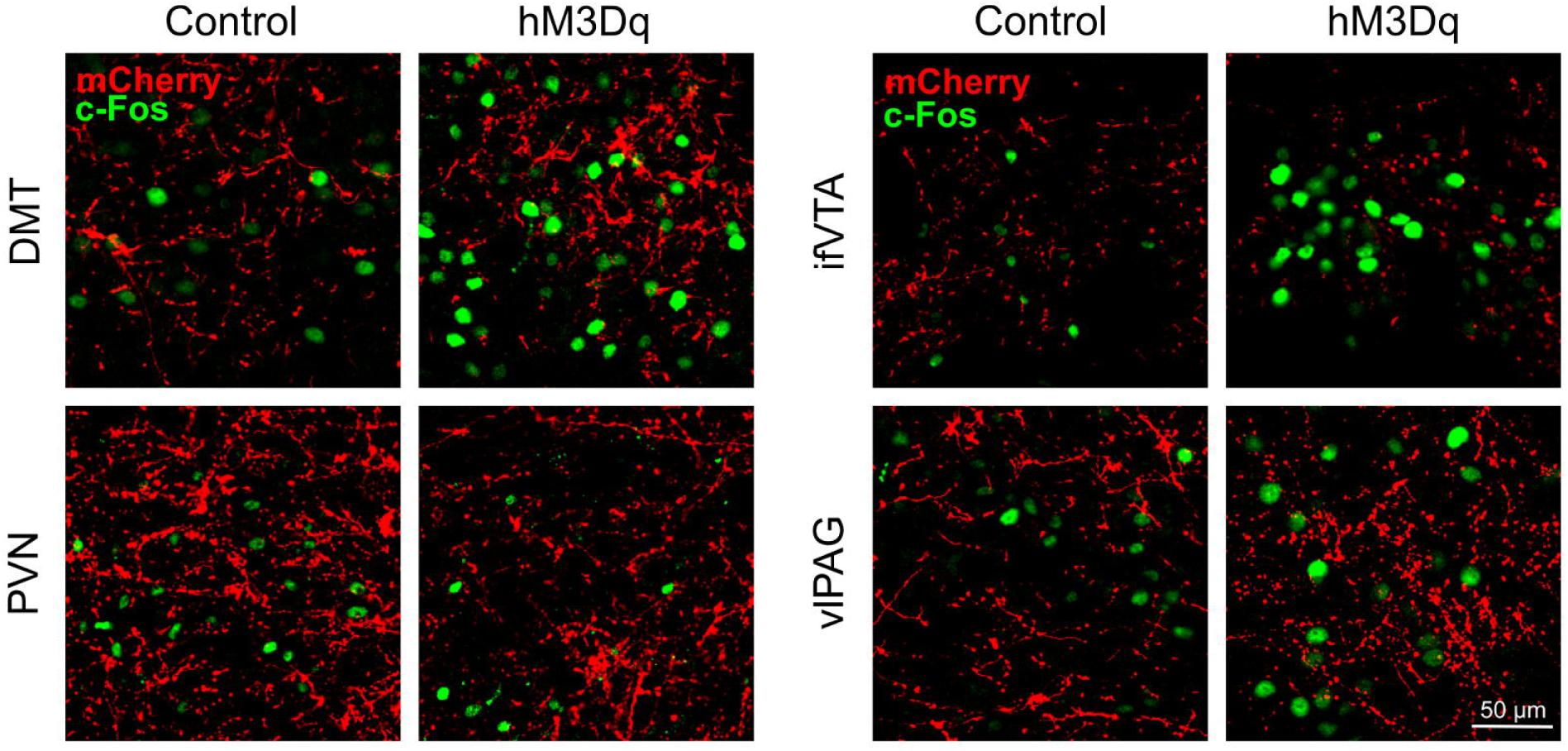
Representative single-plane confocal photomicrographs showing altered c-Fos expression during consolidation in major projection areas of the BNST. *Abbreviations: DMT: dorsal midline thalamus; ifVTA-ventral tegmental area, interfascicular nucleus; PVN-paraventricular nucleus, vlPAG* – *ventrolateral periaqueductal gray.*

**Figure 3 – figure supplement 1.**
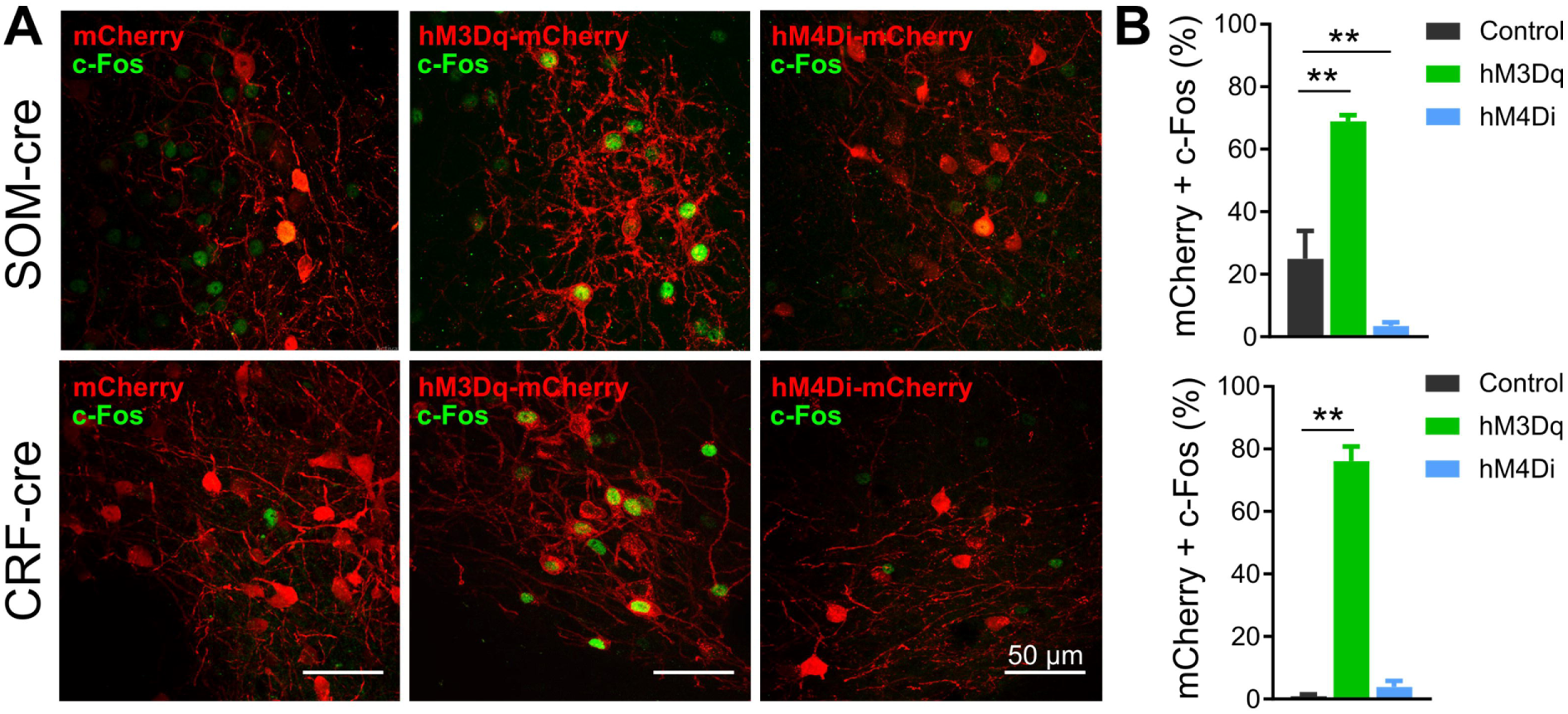
Verification of DREADD mediated modulation of neuronal activity in BNST^SOM^ and BNST^CRF^ neurons. **(A)** Representative z-projection photomicrographs showing c-Fos immunoreactivity in mCherry, hM3Dq-mCherry, and hM4Di-mCherry expressing BNST^SOM^ and BNST^CRF^ neurons after CNO administration. **(B)** Intraperitoneal injection of CNO (1 mg/kg) under baseline (homecage) condition significantly increased c-Fos expression in hM3Dq-mCherry-expressing BNST^SOM^and BNST^CRF^ neurons, while hM4Di expressing BNST^SOM^ neurons showed reduced c-Fos expression (Control: n=3, hM3Dq: n=3 mice). *Asterisks represent main effect of ANOVA: *p<0.05; **p<0.01.*

**Figure 3 – figure supplement 2.**
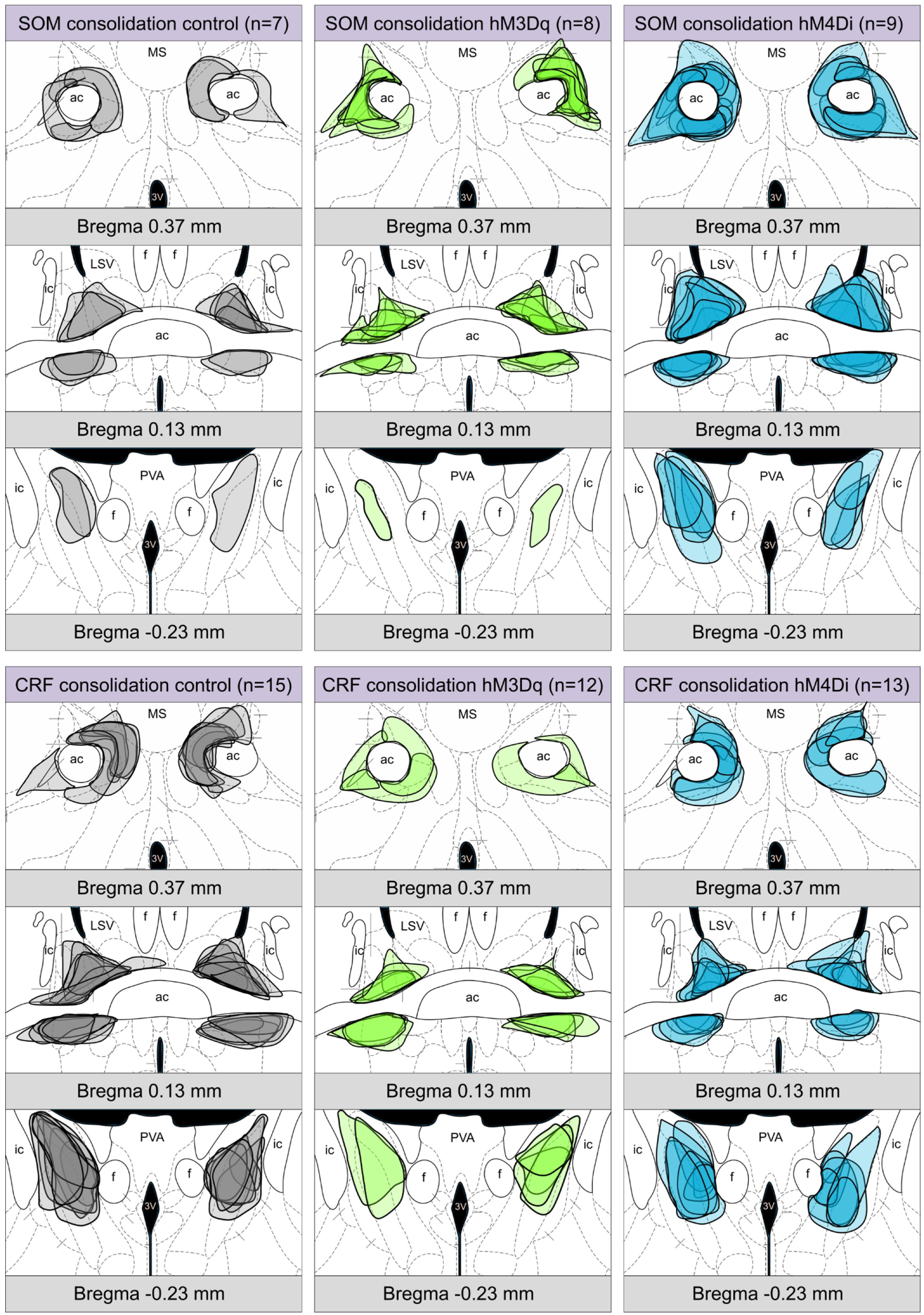
Illustration of AAV-associated mCherry expression in the BNST from control (gray), hM3Dq-mCherry (green), and hM4Di-mCherry (blue) expressing som-ires-cre and crh-ires-cre mice. Diagrams were modified from the Allen Mouse Brain Atlas, Allen Institute for Brain Science (http://mouse.brainmap.org/). *Abbreviations: 3V: third ventricle, ac: anterior commissure, f: fornix, ic: internal capsule, LSV: lateral septal nucleus, ventral part, MS: medial septum, PVA: paraventricular nucleus ofthalamus.*

**Figure 3 – figure supplement 3.**
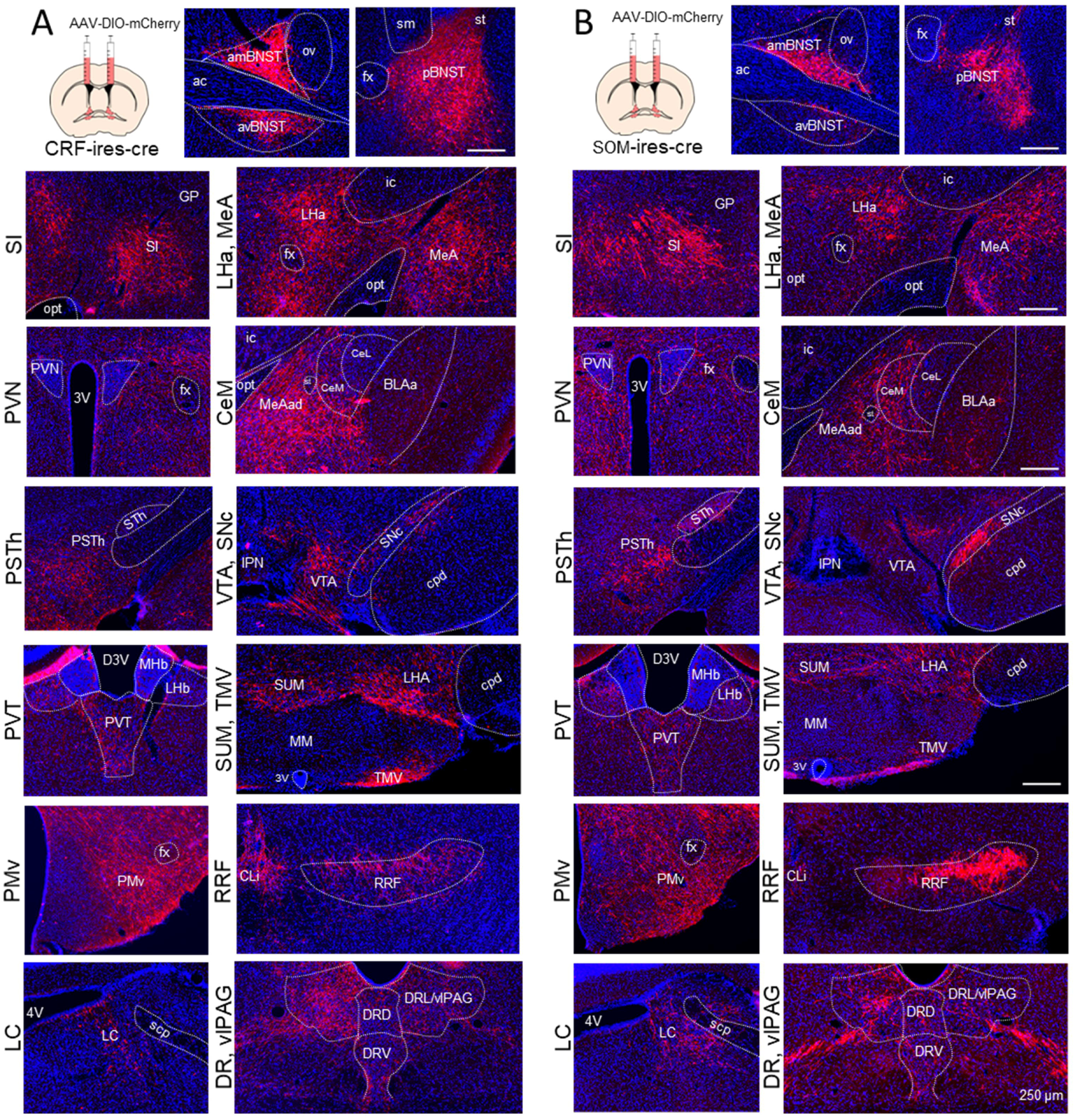
AAV based mapping of major projection areas of BNST^CRF^ and BNST^SOM^ neurons. Representative wide-field fluorescence photomicrographs show a highly similar distribution of BNST^CRF^ and BNST^SOM^ neurons. *Abbreviations: 3V: third ventricle; 4V: fourth ventricle; ac: anterior commissure; amBNST: anteromedial BNST; avBNST: anteroventral BNST; BLAa: basolateral amygdala, anterior part; CeL: central amygdala, lateral part; CeM: central amygdala, medial part; CLi: central linear nucleus raphe; cpd: cerebral peduncle; D3V: dorsal part of the third ventricle; DRD: dorsal raphe, dorsal part; DRL: dorsal raphe, lateral part; DRV: dorsal raphe, ventral part; fc fornix; GP: globus pallidus; ic: internal capsule; IPN: interpeduncular nucleus; LC: locus coeruleus; LHA: lateral hypothalamic area; LHb: lateral habenula; MeA: medial amygdala; MeAad: medial amygdala nucleus, anterodorsal part; MHb: medial habenula; MM: medial mammillary nucleus; opt: optic tract; ov: oval nucleus of the BNST; pBNST: posterior BNST; PMv-ventral premammillary nucleus; PSTh- parasubthalamic nucleus; PVN-paraventricular nucleus; RRF: retrorubral.field; scp: superior cerebellar peduncles; SI: substantia innominate; sm: stria medullaris; SNc-substantia nigra, pars compacta; st: stria terminalis; STh: subthalamic nucleus; SUM: supramammillary nucleus; TMV: tuberomammillary nucleus, ventral part; vlPAG: ventrolateral periaqueductal gray, VTA: ventral tegmental area.*

